# Role of Lactate in the Regulation of Transcriptional Activity of Breast Cancer-Related Genes and Epithelial-to-Mesenchymal Transition Proteins: A Compassion of MCF7 and MDA-MB-231 Cancer Cell Lines

**DOI:** 10.1101/2023.03.23.533060

**Authors:** Inigo San-Millan, Janel L. Martinez, Shivaun Lueke Pickard, Hui Yu, Fred R. Hirsch, Christopher J. Rivard, George A. Brooks

## Abstract

The Warburg Effect is characterized by accelerated glycolytic metabolism and lactate production and under fully aerobic conditions is a hallmark of cancer cells. Recently, we have demonstrated the role of endogenous, glucose-derived lactate as an oncometabolite which regulates gene expression in the estrogen receptor positive (ER+) MCF7 cell line cultivated in glucose media. Presently, with the addition of a triple negative breast cancer (TNBC) cell line, MDA-MB-231, we further confirm the effect of lactate on gene expression patterns and extend results to include lactate effects on protein expression. As well, we report effects of lactate on the expression of E-cadherin and vimentin, proteins associated with epithelial-to-mesenchymal transition (EMT). Endogenous lactate regulates the expression of multiple genes involved in carcinogenesis. In MCF7 cells, lactate increased the expression of EGFR, VEGF, HIF-1a, KRAS, MIF, mTOR, PIK3CA, TP53, and CDK4 as well as decreased the expression of ATM, BRCA1, BRCA2, E2F1, MET, MYC, and RAF mainly after 48h of exposure. On the other hand, in the MDA-MB-231 cell line, lactate increased the expressions of PIK3CA, VEGF, EGFR, mTOR, HIF-1α, ATM, E2F1, TP53 and decreased the expressions of BRCA1, BRCA2, CDK4, CDK6, MET, MIF, MYC, and RAF after 48h of exposure. In response to endogenous lactate, changes in protein expression of representative genes corroborated changes in mRNA expressions. Finally, lactate exposure decreased E-cadherin protein expression in MCF7 cells and increased vimentin expression in MDA-MB-231 cells. Furthermore, by genetically silencing LDHA in MCF7 cells, we show suppression of protein expression of EGFR and HIF-1α, while full protein expression occurred under glucose and glucose + exogenous lactate exposure. Hence, endogenous, glucose-derived lactate, and not glucose, elicited changes in gene and protein expression levels.

In this study, we demonstrate that endogenous lactate produced under aerobic conditions (Warburg Effect) elicits important changes in gene and protein expression in both ER+ and TNBC cell lines. The widespread regulation of multiple genes by lactate and involves those involved in carcinogenesis including DNA repair, cell growth, proliferation, angiogenesis, and metastasis. Furthermore, lactate affected the expression of two relevant EMT biomarkers, E-cadherin and vimentin, which could contribute to the complex process of EMT and a shift towards a more mesenchymal phenotype in the two cancer cell lines studied.

## 1. INTRODUCTION

Described by Otto Warburg a century ago [1] accelerated glucose consumption and lactate production is ubiquitous in solid tumors under fully aerobic conditions. With the discovery of DNA by Watson and Crick in 1953 [2], the landscape of cancer research completely switched from emphasis on metabolism to genetics. At present, while it is recognized that cancer is characterized by mutations of multiple genes involved in cell growth, proliferation, angiogenesis or DNA repair [3], regrettably despite decades of efforts, the cure of cancer through gene science has not been realized. Hence, others and we, have decided to re-open the gates of cancer metabolism studies.

Observations of accelerated glucose uptake and lactate secretion under fully aerobic conditions led Warburg to posit that cancer could be attributed to an injury to the cellular respiratory system (mitochondria) [4], although that idea could not be quite correct [5]. Still, for many decades lactate was considered a “waste” product resulting from lack of oxygen. However, pioneering work and discoveries by George Brooks since the 1980’s demonstrated that lactate is the preferred fuel for most cells, the main gluconeogenic precursor in the body, and a signaling molecule (a “lactormone,” “exerkine and “myokine”) capable of inducing gene expression and ROS generation [6-8]. In concert, Di Zhang et al. have reported recently, the mechanisms by which lactate elicits histone acetylation (“lactylation”). According to them, lactylation is the acetylation of lysine histone residues by lactate, acting as an epigenetic modulator and increasing gene transcription from exposed chromatin in both human and mouse cells [9]. To demonstrate this, Di Zhang and colleagues used isotope-labeled lactate, [3^13^C-lactate] and found a dose-response between lactate levels and lysine lactylation. Furthermore, by tracing labeled glucose [U^13^C-glucose in MCF-7 cells, the same authors showed that lysine lactylation depends on glycolysis as a mechanism to produce lactate. They used several metabolic stressors to promote lactate production through hypoxia and an inhibitor of mitochondrial respiration, rotenone. Further, they suppressed lactate production by inhibiting the reduction of pyruvate to lactate using oxamate. Through a different route, they also suppressed lactate production by using dichloroacetate (DCA) that inhibits pyruvate dehydrogenase kinase (PDK) that inhibits pyruvate dehydrogenase (PDH) that subsequently promotes mitochondrial pyruvate oxidation, thus decreasing lactate production. When lactate production was suppressed, independently of glucose presence, histone lactylation was suppressed. Hence, through their experiments, Zhang and colleagues elegantly demonstrated the mechanisms by which lactate, not glucose, elicited histone lactylation and therefore gene expression [9]. More recently, Jiang et al. have shown that lactate modulates cellular metabolism through histone lactylation-mediated genes in non-small cell lung cancer [10]. As well, Verma and colleagues have also recently demonstrated that tumor-derived lactate can elicit epigenomic reprograming in cancer-associated fibroblasts from pancreatic ductal adenocarcinoma [11]. Moreover, W. Liu and colleagues have recently demonstrated that lactate regulates cell cycle by remodeling the anaphase promoting complex [12].

In 2017, we articulated the “lactagenesis hypothesis” [13] by which we posited that the purpose of accelerated lactate production in tumors was not to support cell bioenergetics [6,14], but rather the role of lactate in tumors was to signal carcinogenesis. Subsequently, in 2019 we demonstrated that lactate functions as an “oncometabolite” and regulates the expression of major genes related to carcinogenesis in MCF7 breast cancer cells [15]. Recognition of the importance of lactate as a hallmark of cancer led several groups to block lactate transport and exchange among and cancer cells and within tumors as a therapeutic approach [16-22].

The central aim of the present study was to investigate the role of endogenous (glucose-derived) lactate as an oncometabolite on important oncogene expression in two representative breast cancer cell lines. Thereby, we sought to extend our findings by studying the representative breast cancer cells lines MCF7, an estrogen-receptor positive (ER+) cell line, and MDA-MB-231, a triple-negative cell line (TNBC). To further confirm the role of lactate in promoting carcinogenesis, we also assessed protein expression for some of the important genes in both breast cancer cells. By using short hairpin RNA (shRNA), we silenced lactate dehydrogenase (LDHA) and LDHA abundance to assess protein expression in MCF7 of two representative genes, EGFR and HIF-1a.

The second aim of our study was to assess the potential role of lactate to regulate the expressions of E-cadherin and vimentin, two proteins that contribute to the epithelial-to-mesenchymal transition (EMT). EMT is a key process in cancer progression characterized by the loss of epithelial, and gain of mesenchymal properties, thus allowing cells to acquire enhanced capacity for motility, migration, invasiveness, and resistance to apoptosis. This increased migratory capacity from a primary tumor site to a secondary, tumor-free site, involves EMT as an important hallmark of metastasis [23,24]. In general, MCF7 cells are characterized by possessing an epithelial phenotype, while MDA-MB-231 display a mesenchymal phenotype [25].

EMT is a complex multifactorial process with four major proteins representative of the process: E-cadherin, N-cadherin, Snail, ZEB1/2 and vimentin. E-cadherin is a transmembrane protein which facilitates intercellular (cell-to-cell) adhesions as well as morphogenesis, limiting cell motility [26]. The decrease of E-cadherin expression disrupts cellular adhesion as well as elicits loss of cellular architecture, facilitating motility and metastasis [27,28]. Furthermore, autocrine signaling reinforces the mesenchymal phenotype and stimulates endothelial growth factor receptor (EGFR), downregulating Ecadherin [29].

Vimentin is a structural protein (type III intermediate filament protein) expressed in mesenchymal cells. Vimentin is attached to the nucleus, endoplasmic reticulum, and mitochondria, anchoring organelles in cytosol [30,31]. Vimentin induces cell migration by inhibiting focal-adhesion proteins [32]. Moreover, it has been shown that vimentin is a key player in metastasis by regulating cell migration and invasion as well as tumor angiogenesis. [33]. Furthermore, it has been shown that TNBC cells possess significantly higher amount of vimentin than ER+ cancer cells [34]. Yongjie Zhang and colleagues demonstrated that lactate promotes EMT in gastric cancer cells by upregulating ZEB2, a transcription factor key in EMT [35]. The approach in this study was to silence LDHA through shRNA in gastric cancer cells. Furthermore, Miranda-Goncalves and colleagues have shown that a 20 mM lactate treatment increased EMT, cell migration and invasion as well as aggressiveness in renal carcinoma cells by downregulating SIRT1 expression [36]. When lactate transporter (MCT) was inhibited, SIRT1 expression increased, and both histone acetylation and cell migration capacity were decreased. Moreover, Li and colleagues claim that lactate is a crucial regulator of EMT, enhancing tumor invasion and metastasis in human lung adenocarcinoma cells [37]. They also showed that lactate decreased E-cadherin expression in a concentration-related manner mainly at a concentration ∼15-20mM lactate. In their study Li and colleagues concluded that the main mechanism by which lactate affects EMT is via increased Snail levels. Finally, Hou et al showed that after LDHA knockdown in lung cancer cells, significant changes in multiple proteins were involved in EMT. They showed increased in E-cadherin and decreased expression of Vimentin, N-cadherin, Snail and ZEB1 [38].

## 2. MATERIALS AND METHODS

### 2.1 Cancer cell lines

MCF7 and MDA-MB-231, both adenocarcinoma breast cancer cell lines were obtained from the ATCC (Manassas, VA) and stored in liquid nitrogen until use. Cells were grown on tissue culture treated dishes (Genessee Scientific, San Diego, CA) in DMEM 1x media with 1g/L glucose (10-014-CV, Corning, Manassas, VA), 10% FBS, and 1X pen/strep. In experiments requiring high glucose media, cells were grown with DMEM 1X media with either 1 or 4.5 g/L glucose (10-013-CV, Corning). In these experiments, the normal 10% FBS was changed to 10% Nu-Serum (#51000, Corning Life Sciences, Glendale, AZ) to reduce the amount of lactate in the initial medium.

Media was also amended with 10 and 20 mM added L-lactate both for 6h and 48h exposures. All cell cultures were grown at 37°C in a laboratory incubator with 5% CO_2_ atmosphere (Forma Isotemp 3530, ThermoFisher, Marietta, OH). Trypsin (0.25% trypsin, 25-053-CI, Corning) was utilized during culture expansion.

### 2.2. QPCR analysis

Cell lines were trypsinized and resuspended in media, followed by low-speed centrifugation to a soft pellet. The cell pellet was then harvested for RNA using the RNeasy Plus Mini Kit (#74134, Qiagen, Hilden, Germany) as per the manufacturer’s protocol. The concentration of purified RNA was determined using a Nanodrop 2000 spectrophotometer (Thermo Scientific, Waltham, MA). cDNA was prepared from 500ng RNA per reaction using the iScript cDNA Synthesis Kit (#1708891, Bio-Rad, Hercules, CA). QPCR was performed using probe-based primer sets and master mix (Prime Time, Integrated DNA Technologies, Coralville, IA) on a CFX96 real time platform (Bio-Rad). Data was evaluated using Maestro software (Bio-Rad) and compared to a standard dilution curve from cell line cDNA. The appropriate house-keeping genes to use for data normalization were determined employing the Human Reference Gene Panel (Bio-Rad). Data was normalized with TBP and HPRT1 analysis.

### 2.3. Cell Protein Isolation and Western Blot Protocol

Western blots for cell line protein expression were performed by harvesting cell cultures from growth plates by removal of spent media, washing plates 3X with cold 1X PBS, followed by addition of cell lysis buffer (#9803, Cell Signaling Technologies, Danvers, MA) amended with AEBSF (1 uM, SBR00015, Sigma) rather than PMSF. Cells were scraped and transferred to a microcentrifuge tube and frozen at -80°C until analyzed. Samples were thawed, sonicated for 5 seconds at 50% power with a pencil type sonicator (model 450, Branson, Danbury, CT). Lysate samples were centrifuged at maximum speed in a microfuge at 4°C for 15 minutes and the soluble protein supernatant removed to a new microcentrifuge tube. Protein concentrations of samples were determined using the BCA protein assay (#23225, Pierce) as per the manufacturers protocol. Protein specimens were aliquoted in 50 uL amounts and frozen at -80°C until analysis. Western blots were loaded at 2.5 to 50 ug total protein per lane in 4-20% gradient pre-cast polyacrylamide gels (#3450033, Bio-Rad). Gels were also loaded with the Dual pre-stained protein standards (#1610374, Bio-Rad) to evaluate protein separation during electrophoresis and transfer to membranes. Gels were transferred to PVDF membranes (#1620177, Bio-Rad) for 2 hours at 250 mAmps followed by blocking in 5% w/v milk in TTBS buffer for 1 hour. Blocked membranes were probed with primary antibodies overnight on a laboratory rocker at 4°C. Membranes were washed 5X with cold TTBS and incubated with anti-Rabbit or anti-mouse secondary antibodies conjugated to HRP (#7074 and #7076, Cell Signaling). After washing, membranes were visualized with ECL reagent (Clarity, #1705060, Bio-Rad) on radiographic film in the dark room. Western blot films were imaged with an Epson 310 scanner (Los Alamitos, CA). Primary antibodies were from Cell Signaling including ATM (#2873), beta Actin (#4970), CDK4 (#12790), E cadherin (#14472), EGFR (#4267), HIF1 alpha (#35169), LDHa (#3582), MET (#8198), MIF (#87501), mTOR (#2983), PIK3CA (#4249), RAS (#67648), and vimentin (#5741), from R&D Systems (Minneapolis, MN) including LDHb (MAB9205), and from Abcam (Cambridge, UK) including Citrate Synthase (AB129095).

### 2.4. Analysis of lactate and glucose concentrations in spent growth media

Lactate in spent growth media was determined employing the L-lactate Assay Kit I from Eton Biosciences (#120001400A, San Diego, CA). Sample lactate analysis was performed in the 96-will plate format with comparison to a standard curve in the range of 30 – 3000uM. Assay plates were measured for the formazan product at 490 nM using a Synergy 2 plate reader (BioTek, Winooski, VT). Net lactate production was determined by subtracting lactate in the initial media from cancer cell’s lactate production at each time point during the experiments.

### 2.5. Seahorse XFe real-time ATP rate assay

For Seahorse experiments, cell lines were maintained in either 1 g/L or 4.5 g/L glucose growth medium for at least 72 hours prior to analysis. Cells were plated at the concentration of about 14,000 cells/well in XFe 96 microplate prior to washing and changing to assay-specific medium sans phenol red and sodium bicarbonate. Brightfield visualization was performed prior to the run utilizing a Cytation Plate Imaging Scanner (BioTek). The real-time measurement of oxygen consumption rate (OCR) and glycolytic extra-cellular acidification rate (ECAR) was performed using the Seahorse XFe 96 Extracellular Flux Analyzer (Agilent Technologies, Santa Clara, CA) with the Seahorse XF real-time ATP rate assay Kit (#103592-100, Agilent Technologies) according to the manufacturer’s protocol. At the conclusion of the assay run, cells are counted via fluorescence in the Cytation reader following exposure to Hoechest 33342 cell-permeable DNA stain (2 μM final, #62249, Thermo Scientific). OCR values were normalized to cell number.

### 2.6. Transfection Protocol for shRNA clones

shRNA vectors for control and anti-LDHA were obtained from the Functional Genomics Core, at the University of Colorado Cancer Center (Aurora, CO). All shRNA vectors were based on the pLKO.1 frame containing a puromycin resistance. The LDHA shRNA vector (TRCN0000164922) was validated for 96% knockdown in HEK293T cells (Sigma, St. Louis, MO). Both vectors were packaged into lentiviral particles prior to transfection. MCF7 cell lines were plated in 24 well plates and on the morning of transfection, growth media was replaced with 0.5 mL Opti-MEM media (31985-070, Gibco, Grand Island, NY) containing 0.5 uL polybrene (TR-1003-G, Sigma) and 10uL to 100 uL of vector/lentiviral preparation. Control transfection was performed utilizing the control pLKO.1 plasmid vector (SCH 002, Function Genomics Core) utilizing the Lipofectamine 3000 reagent (L3000-008, Invitrogen, Waltham, MA) as directed by the manufacturer. For all transfections, following 6 hours of incubation DMEM media was added (1.0 mL) per well for overnight incubation. After an additional 24 hours incubation media was replaced with DMEM 1X 1g/L glucose, 10% FBS, Pen/strep amended with 10 ug/mL puromycin (ant-pr-1, InvivoGen, San Diego, CA). Following daily replacement of media with puromycin, viable growing clones are recovered for evaluation of LDHa protein by Western blot. All evaluations for shRNA clones included comparison to control-vector containing clones.

### 2.7. Statistical analyses

Independent sample t-tests and analyses of variance (ANOVA) were utilized for group-mean comparisons via GraphPad Prism software (version 9.2.1, GraphPad, San Diego, CA).

## 3. RESULTS

### 3.1. Glucose-derived endogenous lactate (The Warburg Effect)

Because lactate was not present in treatment media of either cell line, the lactate present in the spent media was derived from glycolysis (glucose-derived lactate). Glycolysis progressed readily in both cell lines with total and net lactate releases significantly higher after 48h than 6h of incubation (p<0.001) (Fig-1A and 1B, respectively). Notably, lactate release was higher from MDA-MB-231 than from MCF7 cells (69%, p<0.01 at 6 hours, 72%, p<0.0001 at 48 hours) (Fig-1A and Fig-1B). Further, as discussed below, exogenous lactate additions of 10mM and 20mM to incubation media did not affect net lactate production (Fig-1B).

**Figure 1.**
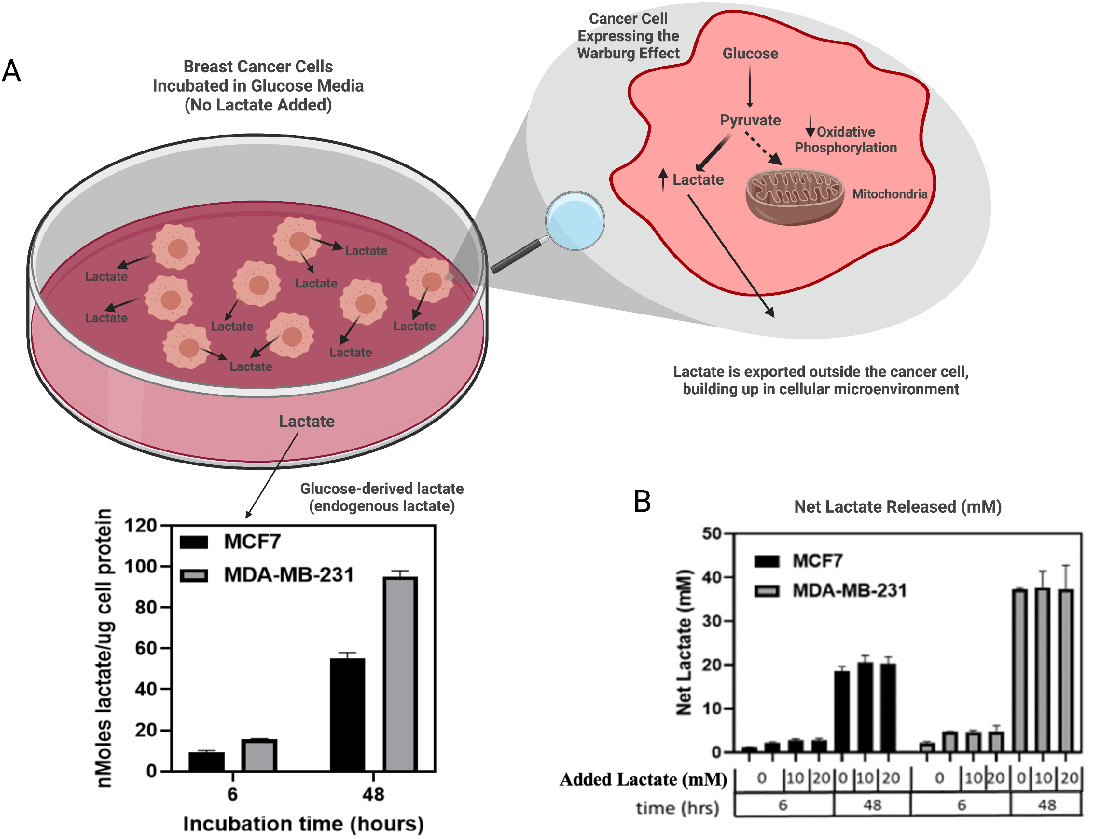
Comparison of lactate production in MCF7 and MDA-MB-231 cell lines (A). Lactate production was measured in the spent media at specific incubation times and normalized for total cell protein. Comparison of net lactate production in spent media as a factor of added lactate, glucose and with increasing incubation time (B). Initial exogenous lactate in growth media was subtracted from lactate determined in the spent media.

The higher glycolytic phenotype of MDA-MB-231 was confirmed via our Seahorse XFe Real-Time ATP Rate analysis. Overall, the total basal ATP production rates of both low-glucose and high-glucose adapted MDA-MB-231 cells were significantly higher than for MCF7 cells (Fig-2A). In MDA-MB-231 cells, more than half of ATP was derived from glycolysis. Once again, by that measure, MDA-MB-231 cells displayed a more glycolytic phenotype than MCF7 cells (Fig-2B).

**Figure 2.**
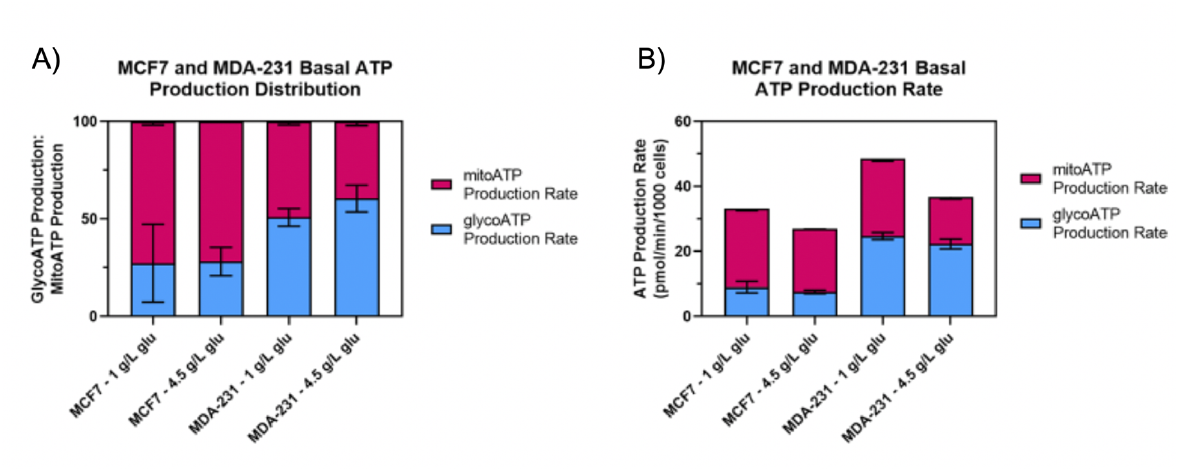
Basal ATP production distribution (A) and total ATP production rate (B) with comparison to breast cancer cell lines and exposure to increasing levels of glucose. Cell lines were maintained in either 1 g/L or 4.5 g/L growth medium for 7 days prior to analysis in test media containing 1.8 g/L glucose.

Consistent with Seahorse analyses, expression showed increased levels of the “lactagenic” enzyme isoform, LDHA, was higher in MDA-MB-231 cells than in MCF7 cells (Fig-3). Conversely, in terms of oxidative phenotype, MCF7 cells showed increased protein expression of the mitochondrial marker citrate synthase (CS) as compared to MDA-MB-231 cells (Fig-3).

**Figure 3.**
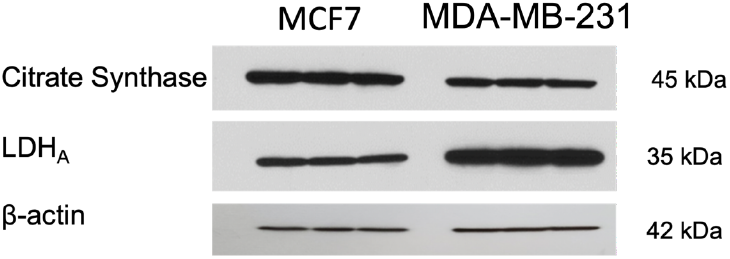
Differences in protein expression in citrate synthase (CS) and Lactate dehydrogenase A (LDHA) between MCF7 and MDA-MB-231 cells representing the higher glycolytic and decreased OXPHOS nature of MDA-MD-231.

### 3.2. Gene expression regulation by lactate

Glycolysis in the cancer cell lines we studied readily resulted in lactate excretion and accumulation (Fig-1). Glucose-derived lactate regulated the expression of multiple representative cancer-related genes in both MCF7 and MDA-MB-231 cell lines (Fig-4). However, this regulation was heterogeneous and varied between cell lines. MCF7 cell lines showed increased expressions of EGFR, KRAS, PIK3CA, mTOR, VEGF, HIF-1α, TP53, MIF, and as well, decreased expressions of BRCA1, BRCA2, ATM, E2F1, CDK6, MET, MYC, and RAF when exposed to endogenously produced lactate, mainly at 48h (Fig-4a).

**Figure 4.**
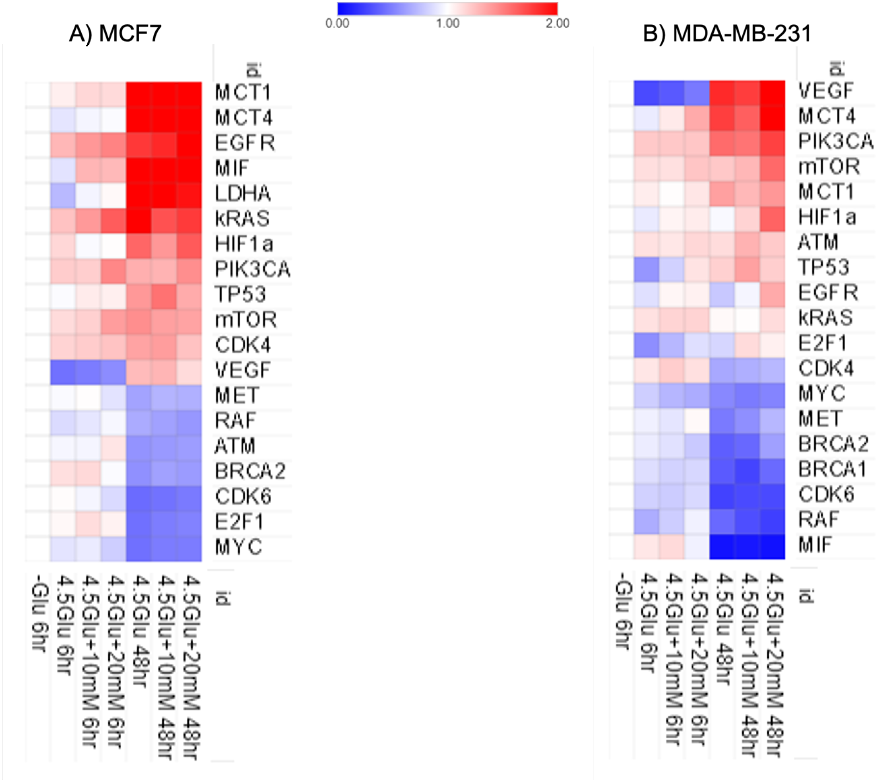
Gene expression after 6 and 48h of endogenous (glucose-derived lactate) and exogenous lactate exposure in both MCF7 (A) and MDA-MB-231 cell lines

In response to glucose-stimulated lactate, MDA-MB-231 cells showed increased expressions of VEGF, HIF-1α, PIK3CA, mTOR E2F1, EGFR and TP53 (Fig-4b). On the other hand, lactate exposure decreased the expression of BRCA1, BRCA2, CDK6, MET, MIF, MYC, RAF, mainly at 48h. However, CDK4 expression increased at 6h, but decreased at 48h (Fig-4b).

The addition of 10mM and 20mM exogenous lactate to glucose-containing media did not increase gene expression compared to endogenously derived lactate and is further explained in Discussion. Noteworthy, lactate increased the expression of monocarboxylate transporters (MCTs) (Fig-4). Genes encoding for MCT1, increased significantly in MCT7 cells, while MCT4 expression increased in both cell lines in response to increased lactate exposure.

### 3.3. Protein expression regulation by lactate

Western blots and their densitometries confirmed that the protein expression patterns of representative genes mirror the expression patterns of the genes identified as having changed from mRNA analyses (Figs-5 and 6). In MCF7 cells, 48h lactate exposure increased protein expression of EGFR, PIK3CA, RAS, HIF-1α, mTOR and CDK4 and decreased the expression of ATM (Fig-5A), confirming data from mRNA expression analysis. In MDA-MB-231 cells, 48h lactate exposure increased protein expression of PIK3CA, mTOR, HIF-1α and ATM, did not affect RAS expression, and decreased the expression of MET and CDK4 (which increased at 6h) (Fig-5B), reflecting mRNA expressions. Like in the case of gene expression, the addition of 10mM and 20mM exogenous lactate did not further increase protein expression compared to endogenous lactate that is further addressed in Discussion.

**Figure 5.**
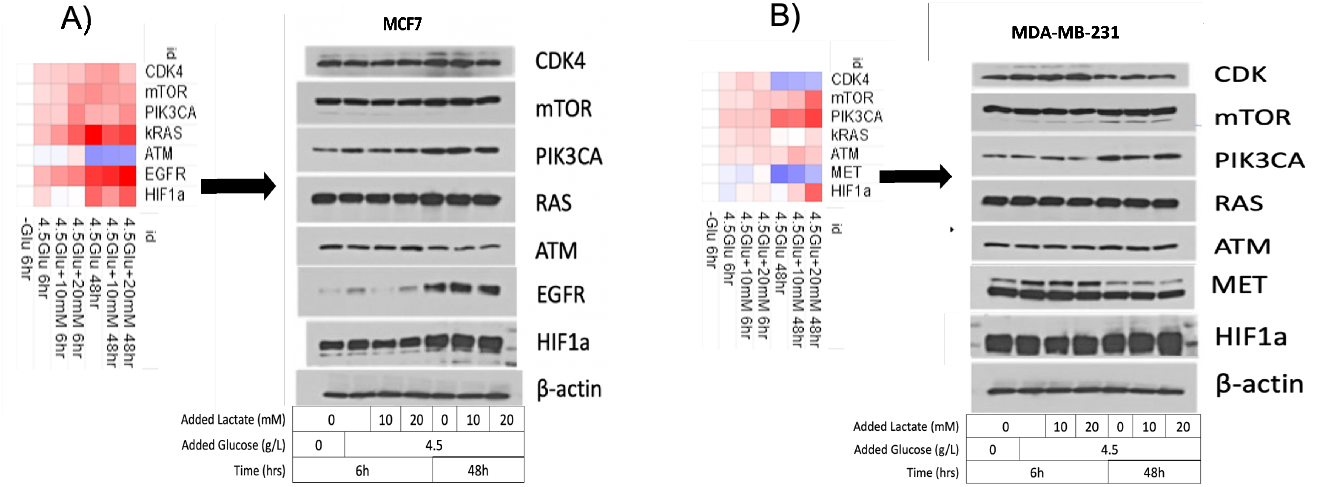
Western blot analysis of protein expression of some representative genes in MCF7 (A) and MDA-MB-231 (B) cell lines following exposure to endogenous lactate (glucose-derived lactate) and added lactate over 6 and 48h of exposure. Specific antibodies are identified in the Materials and Methods section.

**Figure 6.**
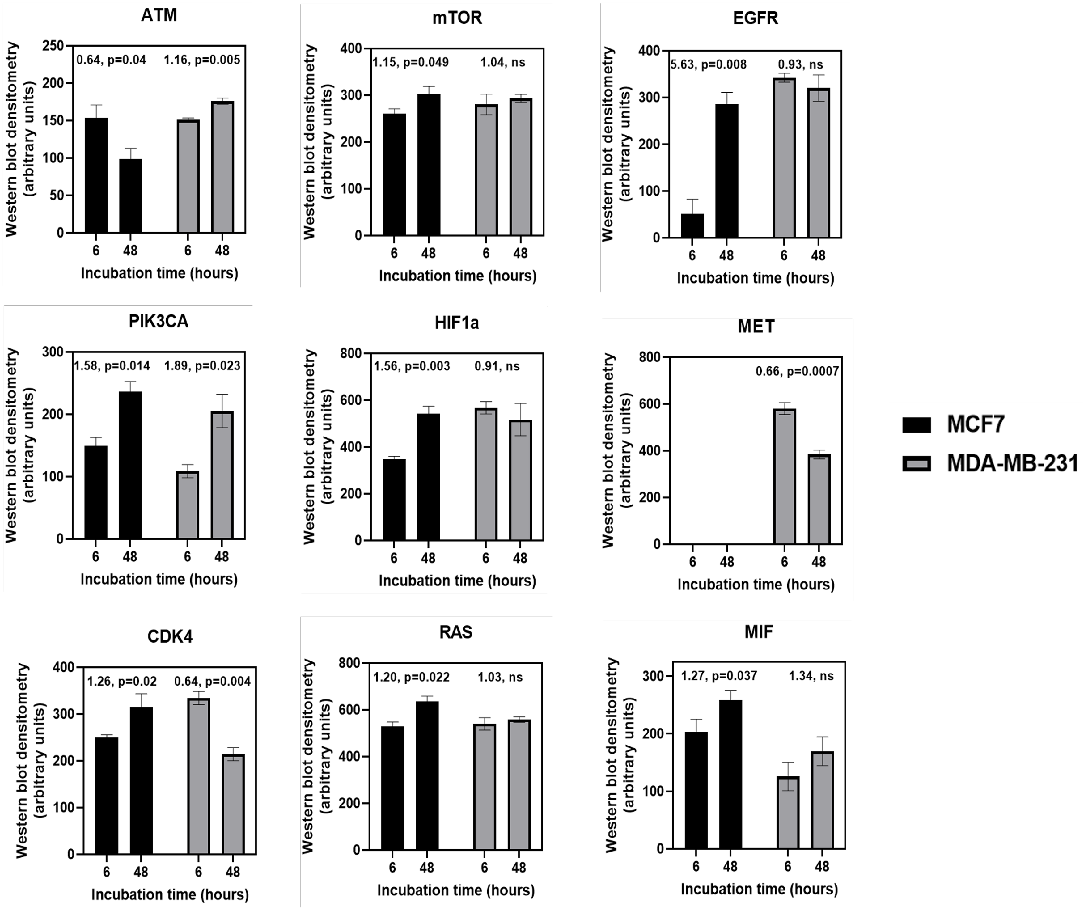
Western blots densitometries of protein expression in MCF7 and MDA-MB-231 cell lines following exposure to endogenous lactate (glucose-derived lactate) and added lactate over 6 and 48h of exposure.

### 3.4. Silencing LDHA in MCF7 cells affects protein expression of EFGR and HIF-1α

In this experiment, utilizing shRNA, we silenced LDHA in MCF7 to assess the protein expression of two representative genes, EFGR and HIF-1α (Fig-7). We observed a complete suppression of the expression of EFGR at both 6 and 48h (Fig-8) as well as a decrease in the protein expression of HIF-1α at 6h and a suppression at 48h when LDHA was silenced. However, under controlled conditions (glucose and glucose + lactate exposures) there were full expressions of both EFGR and HIF-1α (Fig-7). Hence, in this experiment we demonstrate that, glucose-derived lactate as a result of the Warburg Effect and not glucose alone, is responsible for the expression of these two representative genes.

**Figure 7.**
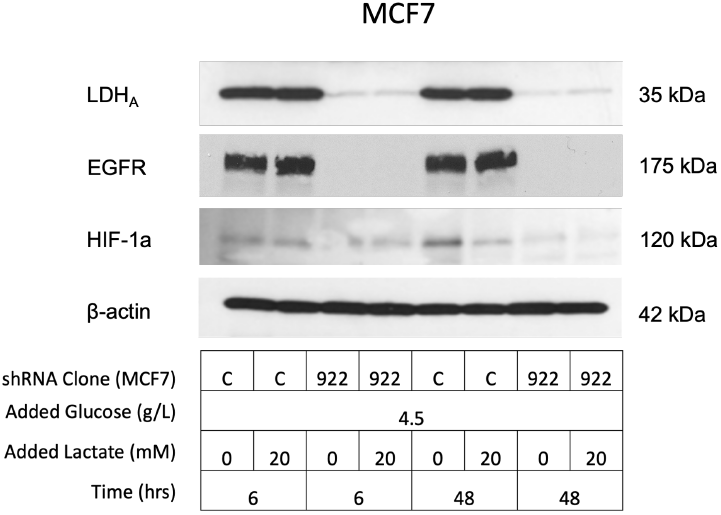
Effects of silencing of LDHA through shRNA on protein expression of EGFR and HIF-1a in MCF7 cells exposed to 4.5g/L of glucose after 6h and 48h. Control shRNA for LDHA (c) shows full LDHA protein expression under control conditions (glucose + glucose +20mM of exogenous lactate) at 6h and 48h. Silencing LDHA trough shRNA (922) shows suppression of EGFR protein expression both at 6h and 48h. Silencing LDHA shows decreased protein expression of HIF-1a at 6h and suppression at 48h.

### 3.5. Possible role of Lactate in the regulation of E-Cadherin and Vimentin

Our results can be interpreted to reveal an important role of lactate as a regulator of E-cadherin and vimentin which are two important regulators of the EMT process in both cancer cell lines. E-cadherin protein expression was decreased by lactate in MCF-7 cells (p=0.058), (Fig-8). Moreover, while vimentin was significantly increased by lactate exposure in MDA-231 cells (p<0.01), we could not detect vimentin in MCF-7 cells (Fig-8).

**Figure 8.**
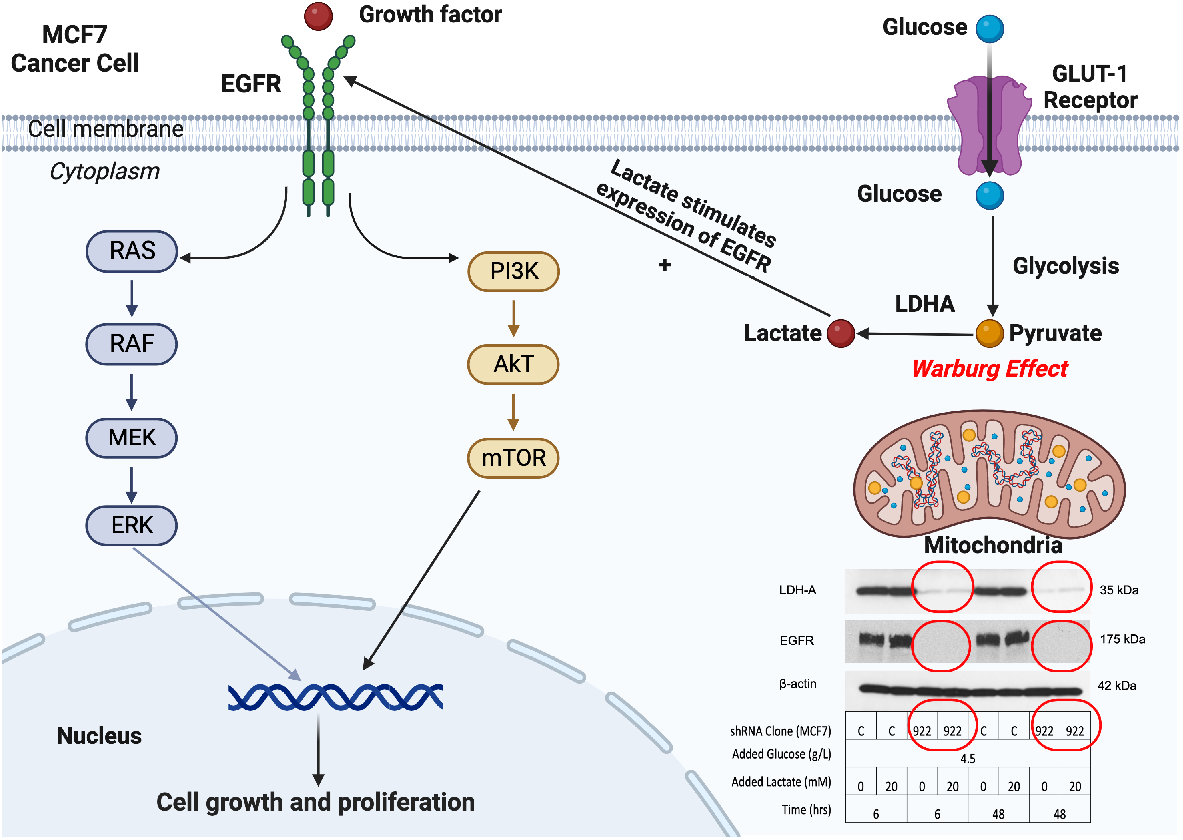
Glucose-derived lactate from the Warburg Effect, stimulates the protein expression of EGRF in MCF7 cancer cells. Silencing LDHA through shRNA (922), both LDHA and EGFR are suppressed (red circles) at both 6 and 48h of exposure to 4.5g/L of glucose.

Like in the case of gene expression experiments, the addition of 10mM and 20mM exogenous lactate did not affect protein expression compared to endogenous lactate production in either cell line (see Discussion).

## 4. Discussion

The role of lactate in regulating the expression of multiple genes is not new. Hashimoto and Brooks previously demonstrated that lactate was able to regulate the expression of 630 genes in multinucleated myotubes and striated fibers-L6 cells [39]. Further, as previously mentioned, Di Zhang et al. demonstrated that lactylation of histone-associated lysine residues serves as an epigenetic modification that directly stimulates gene transcription by lactylation of 28 sites on core histones in human and mouse cells [9]. As well, Jiang et al. have also recently demonstrated that lactate regulates cellular metabolism in non-small lung cancer cells through histone lactylation [10]. Moreover, we recently demonstrated the role of lactate as an oncometabolite, regulating the expression of important cancer-related genes in MCF7 cells [15]. In the present study, we show that lactate is a major regulator of multiple genes and dependent proteins involved in both ER+ and TNBC cancer cells. Furthermore, we show that lactate could play an important role in the regulation of E-cadherin and vimentin which are two important proteins involved in the epithelial-to-mesenchymal transition and vice versa (mesenchymal-to-epithelial transition).

As depicted in Figure 1A, lactate is readily produced by both cancer cell lines studied, with the largest lactate accumulation after 48h of exposure to glucose media. Therefore, the lactate present in media was endogenously produced from glucose, i.e., as a result of the Warburg Effect [1,4]. We also showed this phenomenon in our recent study with MCF7 cells [15]. In the study herein, we observed that MDA-MD-231 cells produce significantly higher quantities of lactate compared to MCF7 cells, with lactate being produced nearly twice as fast as in MCF7 cells per total cellular protein after 48h (p<0.001) (Fig-1). This finding is in line with the higher glycolytic nature of MDA-MD-231 cells reported by others [40-42], which we confirmed via Seahorse cellular respiration analysis (Fig-2). In vivo, the glycolytic phenotype is associated with higher cancer aggressiveness and patient mortality [43-46]. Similar to what is observed in vivo, we found net lactate production at 48h accumulated to 20-40mM in both cell lines (Fig-1B), which is consistent with lactate concentrations found in a variety of tumor microenvironments [47].

To detect the effects of endogenous lactate, we added two different lactate concentrations (10mM and 20mM) to our experimental medium in order to assess whether the addition of exogenous lactate would amplify the expression of multiple genes in a dose-dependent manner vs glucose-derived (endogenous) lactate. Consequently, we concluded that endogenously produced lactate achieved maximum lactate-stimulated responses to genetic and protein expression. The maximum endogenous lactate simulated-response was also true for EMT proteins, E-cadherin and vimentin. Our data shows that endogenous lactate is sufficient to alter genetic and proteomic responses in both cancer cell lines. A main reason for a lack of physiological effect by the additional exogenous lactate in the treatment media could reside in lactate transport kinetics. Lactate is transported into and out of cells by monocarboxylate transporters (MCTs), which are bidirectional, symporters [48,49]. Lactate is transported through cell membranes following proton and monocarboxylate concentration gradients [50,51]. Hence, metabolic activity and physiological processes such as circulatory transport determine the gradients influencing lactate monocarboxylate, mainly lactate release (the physiological Lactate/Pyruvate ranges from 10 to 500, [52,53]) from lactate net producers (driver cells) and consumers (recipient cells) [8]. Thus, cellular lactate efflux as a result of lactate production from glucose (i.e., Warburg Effect) could prevent net influx of exogenous lactate which could probably be the case of our experiments (Figs 4, 5 and 8). Related, is that we observed a significant increase in MCT4 expression in both cell lines at 48h (Fig-4), suggesting a feed-forward stimulus by lactate on MCT4 transporter expression in order to export lactate to the extracellular space where it then becomes a main player in acidosis of the tumor microenvironment (TME) and regulator of multiple TME functions [54,55].

**Figure 9.**
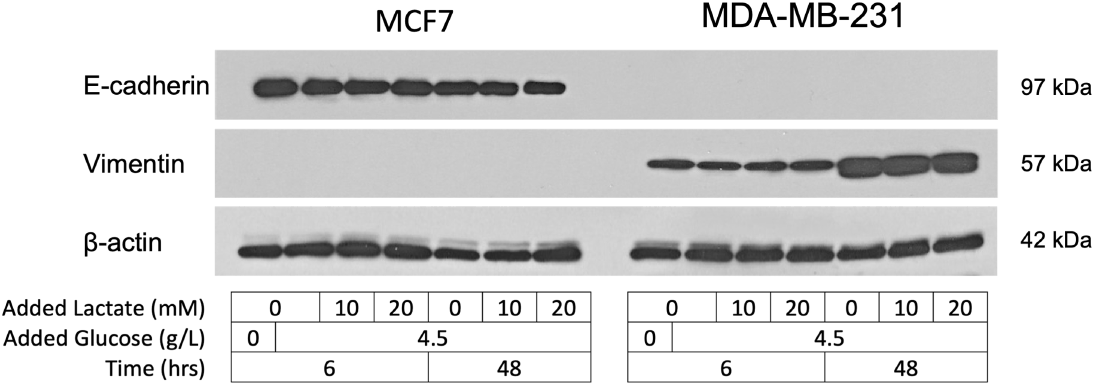
Effect of changes in EMT (Epithelial to Mesenchymal Transition) protein expression in MCF7 and MDA-MB-231 cells with exposure to both glucose-derived lactate and added lactate.

Solid tumors accumulate multiple genetic mutations and in the case of breast cancers, the average number of nonsynonymous mutations is 33, where about 95% of them are single-base substitutions [3]. Within the number of genes mutated, “driver genes” are the key ones conferring cancer cells a selective growth advantage [3,56]. Within these driver genes, it is thought that breast tumors require mutations in three to six driver genes [3]. In both cells studied herein, lactate regulated the expression of multiple important genes involved in all major processes of carcinogenesis including cell proliferation, growth, angiogenesis, metastasis, and tumor suppression (Figs-4 and 5). Most of the genes prevalent in results of our study are considered “driver genes” which are the key genes conferring selective growth advantage to cancer cells, as opposed to untransformed cells [57],.

In MCF7 cells, we observed that lactate elicits significant increases in the expression of genes involved in cell signaling, cell growth, and proliferation like EGFR, PIK3CA, VEGF, mTOR, KRAS, MIF, and CDK4. Lactate also elicited increased expression in genes involved in angiogenesis like HIF1a and VEGF. It is known that lactate stimulates angiogenesis through regulation of the expression of vascular endothelial growth factor (VEGF), a signaling protein that has been known to stimulate the formation and growth of blood vessels [58-61]. Furthermore, Hypoxia-inducible factor 1 (HIF-1) is a master regulator of angiogenesis including the transcription of VEGF [62,63]. According to Semenza, HIF-1 is fundamental for activating the transcription of genes encoding glucose transporters and glycolytic enzymes which elicit metabolic reprogramming from OXPHOS to glycolysis and lactate formation [64]. In our recent study with MCF7 cells [15] and in the study herein, we show that lactate activates the transcription of HIF-1α (Figs-4-5). Hence, there appears to be a reciprocal relationship between HIF-1 and lactate.

Importantly, we found that lactate elicits overexpression of a key tumor suppressor gene, TP53 in both cancer cell lines. Although P53 is typically considered a “tumor suppressor” protein that binds to and repairs damaged DNA or signals for apoptosis in irreparable cells, missense mutations in TP53 elicit mutant p53 proteins with potential gain-of-function properties that stimulate tumor cell proliferation, migration, invasion, survival, chemoresistance, cancer metabolism and/or tissue architecture disruption [65]. Consequently, overexpression of P53 may be associated with increased mortality and poor prognosis [66-68]. Furthermore, we observed that lactate decreased the expression of two other main tumor suppressor factors, BRCA1 and BRCA2, as well as decreased the expression of ATM, a gene involved in DNA repair [69].

In MDA-MB231 cells, lactate increased the expression of genes involved in cell signaling, cell growth, and proliferation including VEGF, PI3CKA, HIF-1α, EGFR, mTOR and CDK4, although the latter only at 6h. As in the case with MCF7 cells, we also observed an increase in the expression of tumor suppressor factor, TP53 which is mutated in ∼80% of TNBCs [70]. The well-known TP53 missense mutation of MDA-MB-231 specifically leads to increased cancer cell survival, migration, and invasiveness. Additionally, like in the case of MCF7, lactate decreased gene expression for BRCA1 and BRCA2.

To corroborate the findings on gene expression obtained through qPCR analysis, we performed Western blots on a representative number of dependent proteins in both cell lines showing lactate’s involvement in protein expression at 6 and 48h of lactate exposure (Fig-5). Overall, it is interesting to observe the differences in gene and protein expression produced by lactate between the two unique breast cancer cell lines. While some key genes and proteins such as PIK3CA, VEGF, mTOR and EGFR were upregulated by endogenous lactate in both cells, there were differences between cell lines in the expression of other genes such as ATM, CDK4, and E2F1. This could be expected given the heterogeneous and different genetic characteristics of both cell lines.

We believe it is noteworthy to observe the role of lactate in regulating epithelial growth factor receptor (EGFR) in MCF7 cells. EGFR is activated via the binding of its specific ligand at the cell surface, epithelial growth factor, and elicits receptor dimerization and phosphorylation, leading to complex signal transduction necessary for cellular proliferation, differentiation, growth, and migration as well as inhibition of apoptosis within the cell. Therapeutically, EGFR is a frontline target in EGFR-subfamily-positive tumors through a variety of approaches including monoclonal antibodies inhibiting both tyrosine kinase receptor and tyrosine kinases [71,72]. These direct inhibitors of EGFR bind to and block the ATP-binding pocket of the receptor, resulting in inhibition of downstream signal transduction. EGFR is a “mother” kinase for two main signaling pathways that are often dysregulated in cancer, which are the RAS/RAF/MEK/ERK and PI3K/Akt/mTOR pathways. It is also noteworthy that PIK3CA, mTOR and KRAS were also affected by lactate in our study herein.

In our study, we showed a significant increase in EGFR protein expression in MCF7 cells in response to endogenous lactate production. This is important in ER+ cancer cells as a decrease in drug response over time and eventual resistance to tamoxifen is a major problem in this largely effective cancer therapy. In these type of ER+ cancers, EGFR overexpression can overrule the signaling inhibition of 17β-estradiol (E_2_) by tamoxifen, increasing carcinogenic signaling through the PI3K/Akt pathway [73]; therefore, anti-EGFR therapy in conjunction with tamoxifen could be preferentially employed in ER+ cancers overexpressing EGFR[74].

To pursue our hypothesis on the role of lactate in carcinogenesis, we decided to silence LDHA through shRNA in order to assess the influence of lactate on the protein expression of two representative genes, EGFR and HIF-1α. We found that silencing LDHA suppressed the protein expression of both EGFR and HIF-1α genes at 48h of exposure while full expression still occurred under control conditions (glucose and glucose + lactate). Hence, in our study herein, we demonstrate that it is lactate, not glucose, what regulates the protein expression of EGFR and HIF-1α.

Finally, regarding EMT, we observed that lactate exposure elicits a decrease in E-cadherin in MCF7 cells that accompanied by a significant increase in EGFR elicited by lactate. That effect could explain some synergies in the decrease of expression of E-cadherin contributing to the loss of epithelial characteristics and positioning lactate as a possible important regulator in this process. On the other hand, MD-MBA-131 cells possess significantly higher amounts of vimentin than MCF7 cells. Lactate exposure significantly increased the protein expression of vimentin at 48h, thus, facilitating a shift to a more mesenchymal, and therefore, metastatic phenotype. Hence, lactate appears to be an important regulator of E-cadherin and vimentin expression in both MCF7 and MD-MBA-131 cells respectively. Therefore, lactate may be an important regulator of EMT in tumor cell lines.

## 5. Conclusions

In summary, endogenous lactate production under fully aerobic conditions, i.e., the Warburg Effect, is capable of eliciting important changes of both gene and protein expression in both MCF7 (ER+) and MDA-MB-231 (TNBC) cell lines. The regulation of multiple genes by lactate is widespread and involves main genes involved in the carcinogenesis process including DNA repair, cell growth, proliferation, apoptosis evasion, angiogenesis, and metastasis. In particular, silencing LDHA in MCF7 cells affects protein expression of EFGR and HIF-1α. Thus, it appears that a key step in carcinogenesis and metastasis is the epithelial-to-mesenchymal transition (EMT) and herein we show that lactate probably plays an important role in the regulation of EMT in breast cancer.

Blocking cellular lactate production as well as the exchange between cancer cells and surrounding stroma cells, affecting the tumor microenvironment, should be a priority in cancer research.

## Author Contributions

Conceptualization, I.S., C.R. and J.M..; methodology, C.R., J.M. and S.L.P.; formal analysis, C.R., J.M, S.L.P, H.Y., G.S. and I.S.; investigation, I.S., C.R., J.M and S.L.P.; resources, C.R., F.H .and I.S..; data curation, C.R., J.M., S.L.P and I.S.; writing—original draft preparation, I.S., C.R. and JM; writing—review and editing, I.S., C.R., J.M. and G.B.; funding acquisition, I.S. All authors have read and agreed to the published version of the manuscript.

## Funding

Funding for this study came from IS-M Cellular Metabolism Laboratory funds and supplementary support from NIH 1 RO1 AG059715-01 to GB.

## Acknowledgments

The authors thank Christina Coughlan, Adriana Solano and Raleigh Jonscher for meaningful discussions of the research results. Use of the Seahorse XFe96 analyzer was provided by Matthew Jackman in the NORC Molecular and Cellular Analytical Core at the University of Colorado Cancer Center.

## Conflicts of Interest

The authors declare that the research was conducted in the absence of any commercial or financial relationships that could be construed as a potential conflict of interest.

## Notes

### Competing Interest Statement

The authors have declared no competing interest.

### Summary of Updates

We have added a new experiment to our study consisting in silencing LDHA in MCF7 cells through shRNA to assess the protein expression of two representative genes, EFGR and HIF-1α (Fig-7). We observed a complete suppression of the expression of EFGR at both 6 and 48h (Fig-8) as well as a decrease in the protein expression of HIF-1α at 6h and a suppression at 48h when LDHA was silenced. However, under controlled conditions (glucose and glucose + lactate exposures) there were full expressions of both EFGR and HIF-1α (Fig-7). Hence, in this experiment we demonstrate that, glucose-derived lactate as a result of the Warburg Effect and not glucose alone, is responsible for the expression of these two representative genes.

